# Physiological regulation of bud burst in grapevine

**DOI:** 10.1101/476879

**Authors:** Santiago Signorelli, Jeremy Shaw, Dina Hermawaty, Zi Wang, Pieter Verboven, John A. Considine, Michael J. Considine

## Abstract

The physiological constraints on bud burst in woody perennials, including the prerequisite for vascular development remain unresolved. Both light and tissue oxygen status have emerged as important cues for vascular development in other systems, however, light requirement appears to be facultative in grapevine, and the information related to the spatial variability of oxygen in buds is unclear. Here, we analysed apoplastic development at early stages of grapevine bud burst and combined molecular modelling with histochemical techniques to determine the pore size of cell walls in grapevine buds. The data demonstrate that quiescent grapevine buds were impermeable to apoplastic dyes (acid fuchsin and eosin Y) until after bud burst was established. The molecular exclusion size was calculated to be 2.1 nm, which would exclude most macromolecules except simple sugars and phytohormones. *In vivo* experiments show that grapevine buds were able to resume growth even following excision from the cane, and that the outer scales of grapevine buds may participate in the biochemical repression of bud burst. Furthermore, we demonstrate that the tissue oxygen partial pressure data correlated well with structural heterogeneity within the bud and differences in tissue density. These data consolidate evidence that the meristematic core becomes rapidly oxygenated during bud burst. Taken together, and when put in the context of earlier studies, these data provide solid evidence that the physiological and biochemical events that initiate bud burst reside within the bud, and question the role of long distance signalling in this developmental transition.

**Highlights:**

- The apoplastic pore size between the grapevine bud and the mother vine is dynamically regulated in the transition to bud burst.
- The molecular exclusion size of the apoplastic connection between the bud and cane is calculated 2.1 nm prior to the initiation of bud burst.
- The structural heterogeneity of the bud explains the spatial variance in tissue oxygen status, and the meristematic core is oxygenated during the initiation of bud burst.
- Long distance maternal signals are not a requirement for bud burst.

## Introduction

Grapevine is the most economically important fruit species worldwide, providing fruit for fresh, dried and processed food and beverage industries, and grown commercially in over 100 countries. The phenology and habit of grapevine is remarkably plastic, displaying strong seasonality and a deciduous habit in temperature regions, while tending towards evergreen in tropical climates (Possingham, 2004). The present understanding of the physiology of these dynamics and the cues that underpin this plasticity is far from complete (Lavee and May, 1997; Considine and Considine, 2016). Improving this knowledge is important to optimise plant productivity, especially in marginal climates or seasons. In particular, an improved knowledge of the physiology of bud burst is fundamental to enable better canopy management and crop forecasting, as the timing and coordination of this event greatly influences flowering, fruitset and ripening.

The axillary buds of several species require light for bud burst or outgrowth (Leduc *et al*., 2014). Several grapevine studies have investigated the influence of low intensity light on bud fruitfulness or shoot physiology, suggesting that it is adapted to a low light environment (Alleweldt and Hofacker, 1975; May *et al.*, 1976; Cartechini and Palliotti, 1995; Petrie and Clingeleffer, 2005; Sánchez and Dokoozlian, 2005). However, there are few reports on the absolute light requirement for bud burst. Although we previously showed that grapevine buds do burst in the absence of light (Meitha *et al.*, 2018), there was evidence of light-responsive gene expression well-before leaf tips emerge through the scale, indicating perception and early resumption of autotrophic capacities (Signorelli *et al.*, 2018). These observations require further physiological elaboration.

Our physiological and molecular data also suggest a developmental role for oxygen (hypoxia) in the initial transition to bud burst (Meitha *et al.*, 2015, 2018), corroborating earlier suggestions from gene expression studies (Or *et al.*, 2000; Ophir *et al.*, 2009; Vergara *et al.*, 2012). More than a third of the widely conserved hypoxia-responsive gene homologues and numerous genes with a hypoxia-responsive promoter element were differentially regulated during the first hours of bud burst (Meitha *et al.*, 2018). Interestingly, we observed that the internal pO_2_ (oxygen partial pressure) minima was peripheral to the meristematic core of the bud following the initiation of bud burst, suggesting an internal source of oxygen (Meitha *et al.*, 2015). However, the spatial resolution of the pO_2_ data was limited in these studies, which we hypothesise reflects the morphological heterogeneity within and between biological replicates. If this is the case, coupling this analysis with x-ray micro-computed tomography (μCT) and tissue density and porosity data should reveal direct correlations and provide a more accurate illustration of the spatial variation and magnitude of pO_2_ within the bud.

An early indicator of the transition to bud burst is ‘sap-flow’; the sudden increase in xylem pressure and the concentration of phytohormones and sugars in xylem sap that precedes bud burst (Skene, 1967; Skene and Antcliff, 1972; Sperry *et al.*, 1987). Shoots are known to provide water and nutrients for bud development, until the bud is photosynthetically active and autotrophic (Michailidis *et al.*, 2018). A role for callose deposition in gating the symplast during the onset and relief of dormancy, metabolically isolating the bud has been illustrated (Aloni and Peterson, 1991, 1997; Aloni *et al.*, 1991; Rinne *et al.*, 2011). Yet, further investigation of this role of callose is required (Beauvieux *et al.*, 2018), as recent reports showed that callose does not explain the changes in connectivity and the molecular size exclusion limit that occurs during development and stress response (Tilsner *et al.*, 2016; Nicolas *et al.*, 2017). In addition, very little is known of the regulation of apoplastic connections during bud burst, which could play a significant role in delivering long range signals from the root or shoot to the bud, given the xylem sap pressure and composition. The evidence and assumptions of recent studies has been that non-cell-autonomous signals that regulate dormancy transitions, such as peptides, are synthesised in embryonic leaves within the bud (Rinne *et al.*, 2011; Paul *et al.*, 2014), however the role of the apoplast in delivering signals cannot be excluded.

Taking these physiological considerations together, we carried out a series of physiological experiments in order to dissect the influences of light, oxygen and apoplastic connection in the regulation of the initial transition to bud burst. We also defined an apoplastic molecular exclusion size for grapevine buds. These studies also provide important chemical modelling data on two major apoplast-mobile dyes, eosin Y and acid fuchsin.

## Materials and Methods

### Plant material and growth under D and DL conditions

Unless otherwise stated, water used throughout the study was Milli-Q^®^ water (MQW, Merck-Millipore, Bayswater, Australia) and chemicals were analytical grade from Sigma-Aldrich (Castle Hill, Australia). Merlot canes containing mature, dormant buds from node 3 to 12 were collected from a vineyard in Margaret River, Australia (34°S, 115°E). The canes were transported to the lab and stored at 4°C until required (no longer than 40 days of storage). Single node cuttings (explants) were prepared as previously described (Meitha *et al.*, 2015, 2018), treated by submersion in hydrogen cyanamide (HC) 1.25 % w/v (Sigma #187364) in water for 30 seconds and then planted on peat. In this respect we consider experimental buds were non-dormant. Explants were grown under dark-light (DL, 12:12h) conditions or in complete darkness (D) and used to perform the experiments described below. The photon flux density for the DL condition was between 200-300 µmol photons.m^−2^⋅s^−1^.

### Bud burst experiments in D and DL

Trays containing 50 explants each were used to quantify the rate of bud burst in D or DL conditions over 52 days. The experiment was performed 4 times using samples collected between late February and early June 2016 (southern hemisphere). The observed response was consistent, with the exception that the time to 50% bud burst decreased as the year progressed, as previously shown in other studies (Or *et al.*, 2002; Parada *et al.*, 2016). The data presented here represent the explants planted on the 2^nd^ May 2017. The explants were treated with HC, or untreated, as indicated in the Results. The trays containing the explants were irrigated with potable tap water every second day to ensure a field capacity of at least 80%.

### CO_2_ release and O_2_ consumption

CO_2_ production was measured as described previously (Meitha *et al.*, 2015, 2018), using 3 pools (n=3) of 8 excised buds each, within an insect respiration chamber (6400-89; Li-COR, Lincoln, NB, USA) attached to a Li-6400XT portable gas exchange system. The measurements were performed in complete darkness, at 23 °C, in CO_2_-controlled air (380 µmol CO_2_ mol^−1^ air) with 100 µmol m^−2^ s^−1^ air flow, at 55–75 % relative humidity.

Oxygen consumption was determined with 5 pools (n=5) of 8 buds each. The buds were placed in a 4 mL micro-respiration chamber (Unisense, Aarhus, Denmark) and the pO_2_ was monitored for at least 10 min using an O_2_ microsensor (Clark-type, Unisense). The calibration of the sensor was performed determining the potential at 0% pO_2_ and atmospheric pO_2_. The of 0% O_2_ condition was achieved flushing N_2_ into the calibration chamber until reaching a constant reading, whereas the atmospheric pO_2_ was determined by flushing air into the chamber and taking into account the temperature of the room. Measurements were performed at 23 °C in complete darkness and keeping the micro-respiration chamber under water to prevent sudden changes in temperature. As a negative control, every day the pO_2_ in an empty micro-respiration chamber was also measured 5 times of 10 min each. The average of the slopes obtained with an empty chamber was subtracted to the measurements of the samples. The volume of the buds was determined using a 25 mL density bottle. This information was necessary to determine for each replicate the final volume of air in the chamber.

Following measurement of the CO_2_ release and O_2_ consumption, the buds were dried in an oven for 72h at 70 °C to determine the dry weight (DW) and the respiration rates were calculated. Note that RQ values cannot be deducted from this data because the conditions of measurements are different for CO_2_ and O_2_.

### Internal pO_2_ profiles

The internal O_2_ (pO_2_) was measured using an O_2_ microsensor (Clark-type, Unisense) with a 25 µm tip as previously described (Meitha *et al.*, 2015, 2018). The calibration of the sensor and measurements were done using the Sensortrace Suite software. For calibration of 0 kPa O_2_, the sensor was flushed with N_2_ until stabilization, and for normal pO_2_ concentration, a fish pump was used to flush the sensor with atmospheric air. The buds were removed from the cane and stood on a flat surface and then the electrode was electronically inserted from the top to reach the centre of the meristem as described in Meitha *et al*. (2015). The depth of the path changed depending on the size of the bud, but it was usually between 2900-3700 µm with steps of 36 µm and 3 measurements per step. The profiles were performed in at least 8 buds for each condition and time (n=8).

### Apoplastic connection

Both, acid fuchsin and eosin Y solutions were prepared dissolving 0.5 g of the respective dye in 0.01 M phosphate buffer pH 6.0. The solutions were filtered through GFA filter paper and then through a 0.2 µm membrane filter. Explants were sampled from growth conditions at the corresponding times and cut 1.5 cm beneath the bud and immediately transferred to 25 mL plastic tubes containing 2 mL of eosin Y or acid fuchsin solutions. Unless otherwise stated the incubation time in the solution was 24h. A razor blade was used to make a longitudinal cut through the centre of the buds, and the images were taken using a camera, coupled to a magnifying glass. At least 3 replicates were used per dye per time (n=3).

### Vascular connection and porosity by micro-computed tomography

Grapevine canes collected on the 4^th^ April 2016, were sent to Belgium via World Courier and kept at a constant 4°C. The explants were planted on the 23^rd^ May 2016 in D and DL as was done for bud burst analysis. Three buds of each condition (0h, 48h, 72h, 168h each D and DL) were incubated overnight with CsI 10% at 4 °C. 3D imaging using micro-computed tomography (µCT) was performed on each bud. µCT was performed with a Phoenix Nanotom (General Electric, Heidelberg, Germany) before and after incubation. For scanning, buds were mounted on the rotation stage by means of a Parafilm wrap. 2400 projection images per scan were taken with 0.15° angular steps for a full 360° rotation. Capture time for each image was 500 ms. Settings were 55kV/182uA for control samples, and 60kV/167uA for CsI samples. Image pixel resolution was 2.50 to 3.25 µm depending on bud dimensions. Slice reconstruction was performed by Octopus Reconstruction version 8.9.0.9 (XRE, Ghent, Belgium) using the filtered back projection method.

3D image rendering and quantitative image processing was conducted in Avizo 9.6 (ThermoFisher Scientific, Bordeaux, France). First, the bud volume was masked from the background and the Parafilm wrap used for scanning. This was achieved by applying a global threshold on the grey scale images complemented with erosion, dilation and filling operations on orthogonal image slices and the 3D volume. Second, the pixels inside each masked bud image were assigned to air or bud tissue using a simple grey scale threshold. Pixel values lower than or equal to 60 (on a 0 to 255 scale) were assigned to air, according to the greyscale range of the background. Pixel values higher than 60 were assigned to the tissue. Finally, different subsamples (with a representative volume larger than 100 by 100 by 100 pixels) of the different parts of the bud (outer and inner scales, trichomes, base) were taken from the 3D image to calculate local porosity. Porosity was defined as the proportion of total volume of air to the total volume of the subsample. Porosity values of tissues were averages of 3 buds per condition.

### Molecular modelling

The geometrical structure of associated and dissociated forms of rhodamine green, acid fuchsin and eosin Y was fully optimized in aqueous solution at the B3LYP/6-31G(d,p) level integrated with the IEF-PCM polarizable continuum model without imposing symmetry restrictions and using solute cavities adapted to the molecular shape and constructed with Bondi radii. The nature of each minimum was verified by inspection of the eigenvalues of the analytic Hessian in aqueous solution. Optimizations were performed at 298.15 K in the Gaussian09 program. All the calculations were performed using Gaussian09, rev. A.1 (Frisch *et al.*, 2009). To visualise the molecules, animate the vibrational modes and represent the electron density and the charges, the program Gaussview 9.0 was used. The solvent-accessible surfaces (or Lee-Richards surfaces) were determined using Jmol (The Jmol Team, 2007).

### Effect of scales on bud burst

To determine the effect of the scales in bud burst the outer scale of the buds was removed using razor blades and forceps. These experiments were performed in the absence of HC. At least two replicates of 30 buds each were used for buds with and without scale in the different conditions (DL and D; n=2).

### Bud porosity and humidity

Five sets of five buds each (n=5) were used to determine moisture content and porosity. Each bud was dissected to separate the scale, trichomes and green tissues. The porosity and moisture were measured for each tissue. Fresh weight was recorder before measuring porosity. After measuring the porosity, the tissues were desiccated during 96h in an oven at 70 °C and the dry weight determined. The moisture content was determined as follows:

Moisture (%) = 100 * (FW – DW) / FW

To determine the porosity the volume of the tissues and the volume of air in the tissues was determined using a density bottle (Thomson *et al.*, 1990). The porosity was measured as follows:

Porosity (%) = 100 * Volume of gas in the tissue / Volume of tissue

### Bud burst on isolated buds

Following treatment with HC 1.25%, 20 buds were peeled by removing the outer scale, and an equal number left intact. Then the buds were excised from the cane and placed in a solid Murashige Skoog (MS) medium, agar 0.8%, containing Plant Preservative Mixture (PPM) at 0.2%. The buds were grown in the plate for 12 days under dark-light (DL, 12:12h) conditions. To evaluate growth, longitudinal sections were performed and they were observed under a magnifying glass. Digital images were taken with the camera coupled to the magnifying glass and with a ruler close to the buds as reference. The distances (height and width of bud and primary bud) were evaluated digitally.

### Effect of oxygen on bud burst

Two trays containing 60 explants were placed in a custom gas-tight transparent chamber connected to an O_2_ cylinder (99% O_2_). For the hyperoxia treatment, the chamber was flushed to saturation with 99% O_2_ for 30 min each day for 15 days. Before doing the O_2_ treatment every day the chambers were open to renew the air and remove the moisture, and when necessary they were irrigated. A similar procedure was performed in the control chambers but flushing the chambers with air instead of O_2_. In addition, the hyperoxia treatment was performed with trays containing intact and peeled explants (outer scales removed).

### Electrode pathway by micro-computed tomography

The buds used to determine the internal pO_2_ were preserved in FPA solution (10% (v/v) 37% formalin; 5% (v/v) propionic acid; 50% (v/v) 70% ethanol; 35% (v/v) DI water) at 4 °C. The samples were subsequently incubated overnight in 1% iodine +2% potassium iodide solution (IKI) in PBS prior to µCT scanning. Buds were then placed in a 5 mm diameter sealed plastic straw in PBS and scanned at 40 kV and 66 µA using a µCT system (Versa 520 XRM, Zeiss) running Scout and Scan software (v10.6.2005.12038, Zeiss). A total of 2501 projections were collected over 360°, each with a 5 s exposure. 2x binning was used to achieve a suitable signal to noise ratio and 0.4x optical magnification was used to achieve an isotropic voxel resolution of 7.9 µm. An LE1 source filter was also applied. Raw data were reconstructed using XMReconstructor software (v10.7.3679.13921, Zeiss) following a standard center shift and beam hardening correction. The standard 0.7 kernel size recon filter setting was also used. Avizo (v8.1.1, FEI) software was used to obtain orthogonal slices through the data in the same plane as the electrode pathway. Images were then taken into Adobe Photoshop Elements 15 where the the pO_2_ profiles were mapped onto the electrode pathway. Pixel density was determined using the Line probe tool of Avizo 8.1, as a measure of tissue density. In particular, two lines were used, one just above the path and the other just below the path. The average of the intensities was used to estimate the density of tissue that the probe penetrated.

## Results

### Physiological experiments on the roles of light and oxygen during bud burst

The influence of light on the rate of bud burst was evaluated in the presence or absence of hydrogen cyanamide (HC), a commonly used agent to synchronise bud burst. The data demonstrate an interaction between light and HC, whereby the influence of light was greater in the absence of HC (Fig. 1A,B). HC alone had a considerable effect of increasing the rate of bud burst, accelerating the rate to achieve 50% bud burst from *c*. 28 to 15 days (in light). In the absence of both light and HC, bud burst did not reach 50% within the experimental timeframe. However, light was not an obligate requirement for bud burst. For subsequent experiments we chose to work with HC-treated buds because bud emergence was more predictable than in untreated buds. Where exceptions occurred, they were noted.

**Figure 1.**
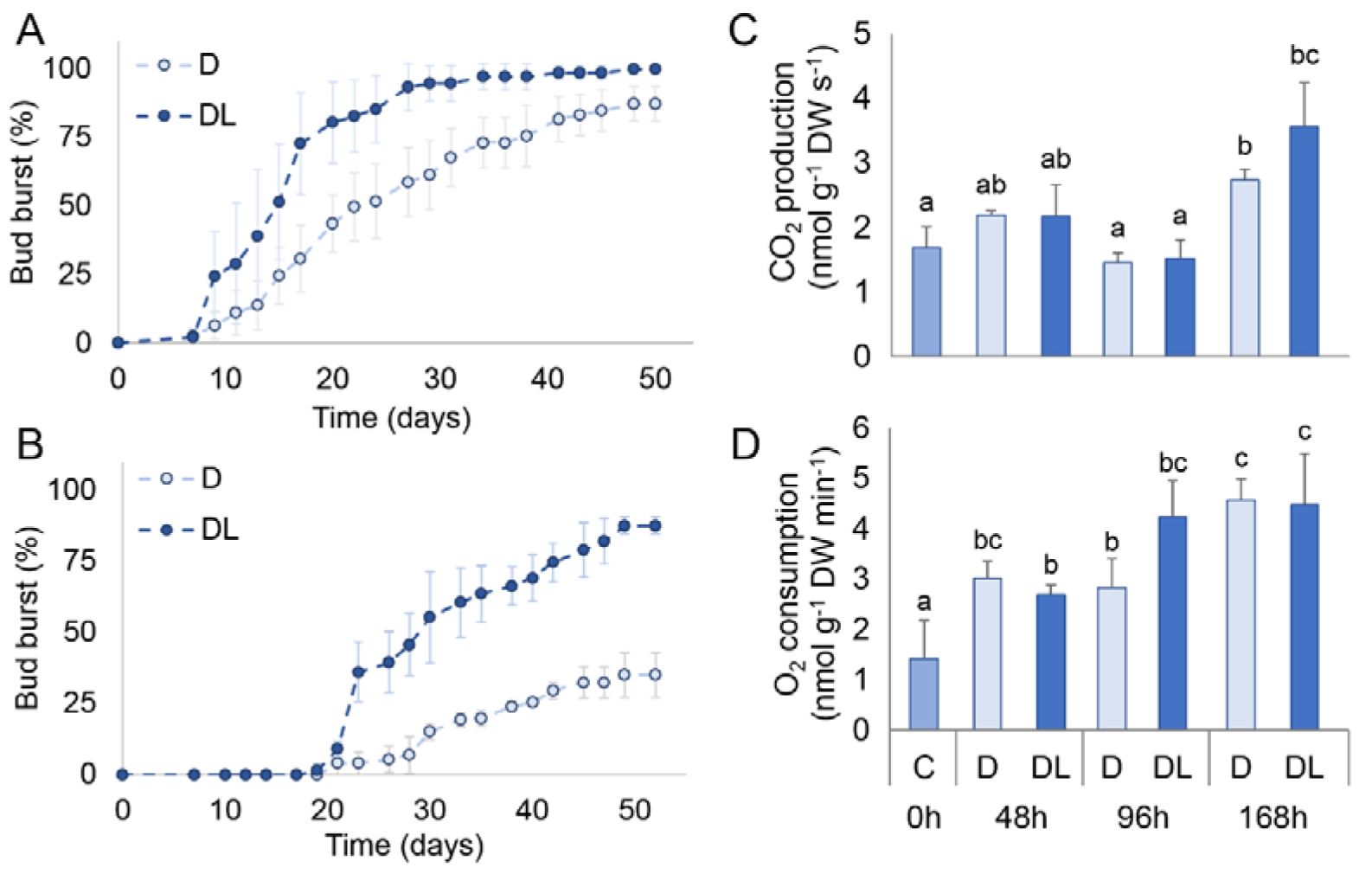
Effect of light on bud burst and bud respiration. Bud burst percentages of grapevine buds treated with HC 1.25% (**A**) and untreated (**B**), grown under dark-light (DL, closed circles) and darkness (D, open circles) conditions. The bars indicate standard deviation, n=3. Respiration determined by CO_2_ release (**C**) and O_2_ consumption (**D**). Different letters indicate significant differences against the respect to control (0h) using a Tukey comparison (n=4, *p*< 0.05).

Bud respiration was determined in the presence and absence of light (DL, D) as CO_2_ production and O_2_ consumption during the first week of growth (0, 48, 96 and 168h). In both light conditions at 168h, the CO_2_ release and O_2_ consumption were greater than at 0h (Fig. 1C,D), indicating the resumption of metabolic activity irrespective of the presence of light. Although minor differences in gas exchange were observed between the DL and D conditions, these were not significant and not as apparent as previously observed (Meitha *et al.*, 2018).

Assuming that the outer scales of grapevine buds might act as a barrier to O_2_ diffusion, we analysed the effect of removing the outer scales on bud burst. Peeled buds were able to burst earlier than unpeeled buds, as previously observed in var. Zinfandel (Fig. 2A; Iwasaki and Weaver, 1977), suggesting that the scales of grapevine buds delay bud burst. Initially we considered two possible explanations for the acceleration of the bud burst: an increased incidence of light may stimulate bud burst; or, an improved gas exchange may promote oxygenation, relieving a limitation to respiration. To identify whether light was playing a role, we performed the same experiment in absence of light (D), showing a delay in bud burst relative to light condition (DL) of both control and peeled buds (Fig. 2A,B). The difference however between peeled and control was not suppressed by the absence of light. This refutes the argument that light incidence was primarily responsible for the acceleration of bud burst in peeled buds. We also tested the rate of bud burst of intact buds in a hyperoxic environment, but found no difference to normoxic conditions, and the difference between peeled and control was not suppressed (Fig. 2C). Together this suggests that the inhibitory effect of the outer scale on growth is other than reducing light or oxygen perception. This is considered further in the discussion section.

**Figure 2.**
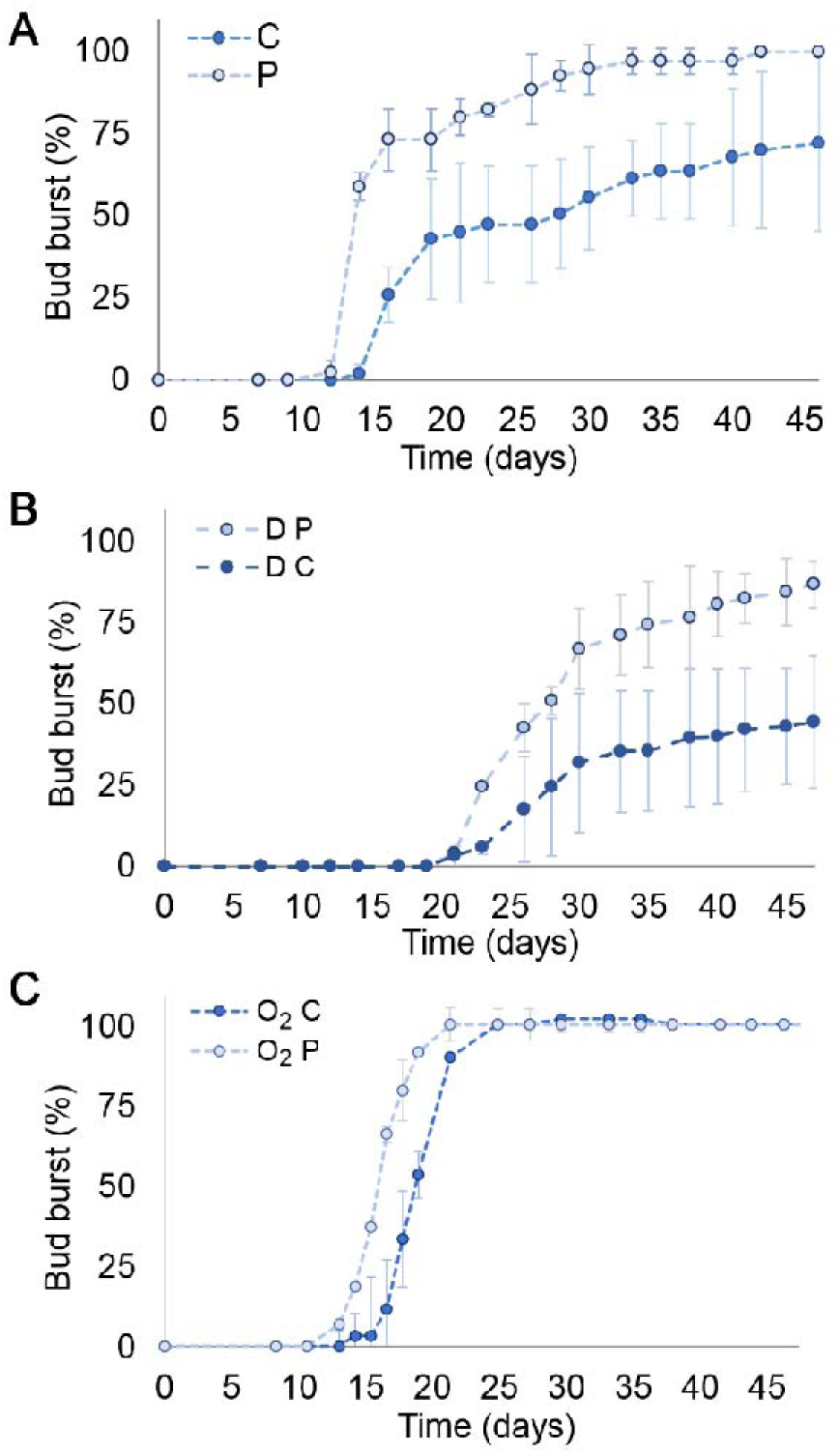
Effect of scales on bud burst. (**A**) C, control bud. P, peeled bud; (**B**) D C, darkness treatment in control buds. D P, darkness treatment in peeled buds. (**C**) O_2_ C, hyperoxia treatment in control buds. O_2_ P, hyperoxia treatment in peeled buds. The vertical bars indicate standard deviation.

### The role of vascular development during bud burst and the effect of light

We then investigated the vascular development of the intact buds visually and by micro-computed tomography (µCT), to address whether light influenced the resumption of vascular transport. From the μCT data, vascular development was apparent over the time course however the contrast agent was transported even at 0h) demonstrating that the vascular tissue is already functional, at least, to transport water and minerals (Fig. 3A and Movie S1 show the uptake of the contrast agent in a 3D bud structure). The contrast agent used here, was determined to be ideal for marking vascular tissue (Wang *et al.*, 2017). On the grounds that the symplast was shown to be regulated during the dormancy transitions in poplar (Rinne *et al.*, 2011), we investigated whether the apoplast is also regulated during the resumption of bud growth, using apoplastic dyes which are larger than the contrast agent used for µCT and more reflective of macromolecules transported in the phloem and xylem. Here, dye movement was not evident until after the swollen stage (168h, 7 days), when bud break was already initiated (Fig. 3B,C). The ability of the apoplast to transport the dyes was observed earlier in the DL condition, despite the fact that D-grown buds had reached a similar degree of bud swell over the same time (see Fig. S1 and S2). This demonstrates that the aperture of the apoplast between the bud and the cane is rapidly regulated during the initation of bud burst and likely to restrict uptake to water and small molecules until after the initiation. It also suggests that the difference in the rates of bud burst observed between D-and DL-grown buds results, in part, from more rapid apoplastic development in the DL condition (Fig. 1A). In addition, we observed that the uptake of acid fuchsin was more rapid and extensive than of eosin Y.

**Figure 3.**
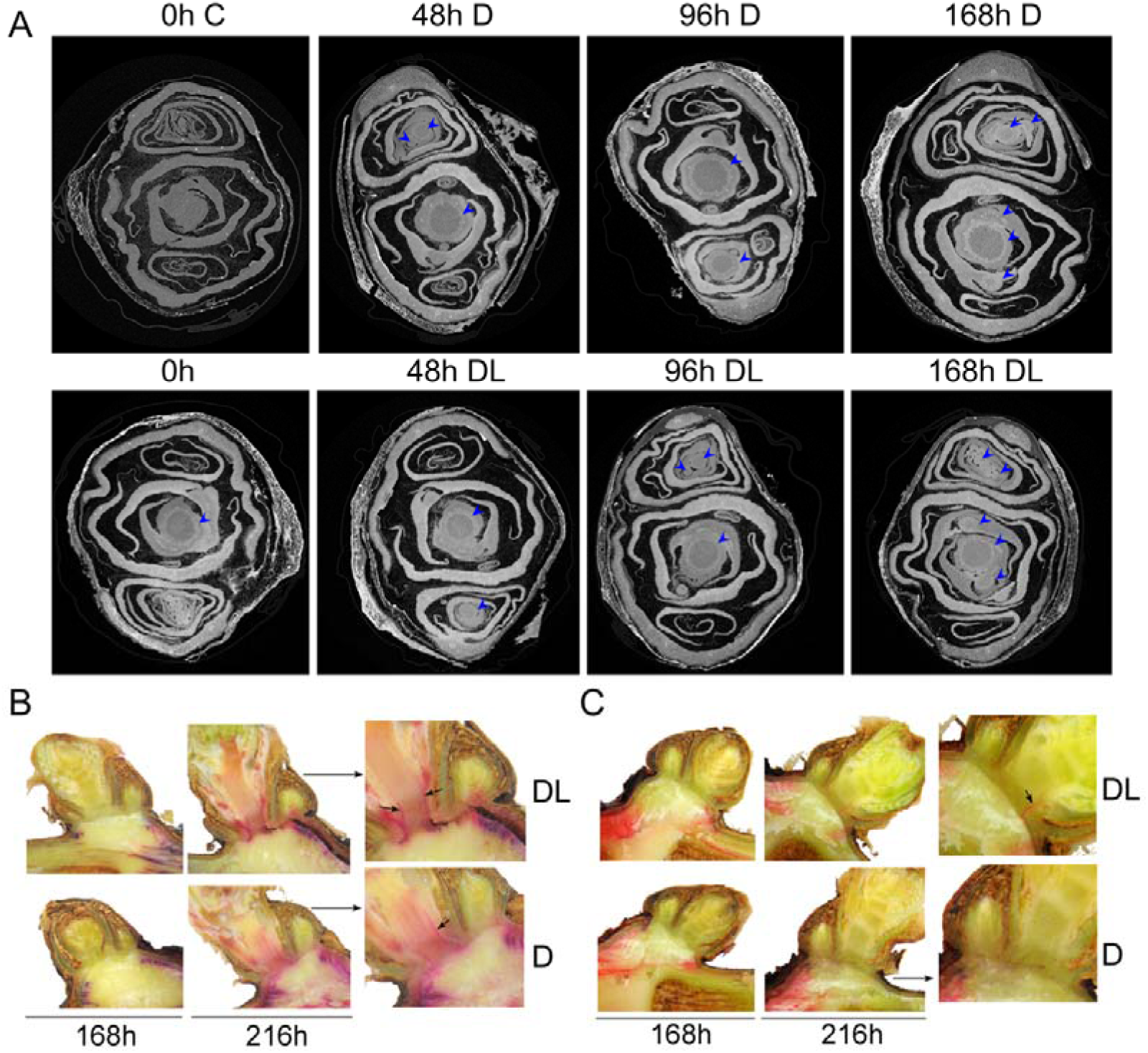
Effect of light on vascular development. Micro-computed tomographies of buds (**A**). 0h C refers to 0h buds untreated with contrast agent caesium iodide (CsI). All the other buds were treated with contrast agent in order to visualize the vascular tissue (observed as rings and indicated with blue arrows); Apoplastic connectivity evidenced by acid fuchsin (**B**) and eosin Y (**C**) staining. Arrows in the magnified image indicate Eosin Y staining.

Considerable evidence shows that the xylem pressure builds and the sap becomes enriched with simple sugars and phytohormones, particularly cytokinins, in the weeks prior to bud burst (Skene and Antcliff, 1972; Sperry *et al.*, 1987; Maurel *et al.*, 2004; Bonhomme *et al.*, 2010). The data described above indicate these smaller molecules would be capable of being delivered to the meristematic cells of the bud via the apoplast, but not larger oligosaccharides, peptides or macromolecules which may play signalling roles. Thus we performed a bud burst experiment isolating the buds from the cane and planting them on Petri dishes containing MS-agar-PPM. Considering the beneficial effect of removing the scales on bud burst we used intact buds and peeled buds, where the scale was carefully removed. We observed that excised buds from late September (2 weeks prior to natural bud burst) could initiate bud burst, since the primary buds were already swollen at 12 days in peeled buds (Fig. S3). Intact buds were also swollen at 12 days (not statistically significant from control). Despite the swelling, the growth was limited and the buds were unable to sustain leaf emergence (bud burst *sensu stricto*), possibly due to nitrogen, phosphorus or other mineral deficiency, otherwise supplied through the transpiration stream from the cane.

Since the transpiration rate in a later stage of bud burst should be faster than in an earlier stage, it is plausible that the greater uptake of dyes at later stages is due to the greater transpiration, rather than a change in apoplastic connectivity. Thus, we performed an experiment comparing bursting buds with 0h buds that had been peeled in order to increase the dehydration of the bud, and therefore the uptake of the solution. Our results demonstrate that in an intact bursting explant (216h), the dye penetrated to the top of the bud within 3h of incubation (Fig. S4A). Nevertheless, in a 0h peeled bud, even after 72h of incubation, the dye could not penetrate the buds (Fig. S4A). This demonstrates that the observations of Fig. 3B,C are methodologically supported, and that prior to bud burst, bulk flow driven by the xylem pressure, or diffusion would still be restricted.

Following this series of experiments, we conclude that neither light incidence, nor long distance signals arising from the mother vine are essential for the resumption of organogenesis leading to bud burst.

### Determining the tissue-specific oxygen levels in a perennial bud

In order to further investigate the roles of light and O_2_ on the course and coordination of bud burst, we assessed the internal tissue oxygen status (pO_2_) through bud burst in the DL and D buds. These data were consistent with our previous studies, showing progressive spatio-temporal oxygenation during bud burst (Fig. S5; Meitha *et al*., 2015; 2018). While a slight decrease in the pO_2_ over the first 48h and a marginal increase after 48h was observed, the pO_2_ differences we had previously observed between the DL and D conditions (Meitha *et al*., 2018) were not as apparent. These data were measured in complete darkness (assayed condition), however, the presence of chlorophyll (Meitha *et al.*, 2018) and light-responsive gene expression (Signorelli *et al.*, 2018) during the initiation of bud burst suggests sufficient light does penetrate and promote light signalling and may enable photosynthesis. Thus, we re-examined the pO_2_ profiles in the presence of light (assayed condition), with the expectation that the presence of light promotes internal oxygenation via photosynthesis and may be related to the more rapid bud burst observed when light is present in the growth environment. This experiment was performed only in 168h buds, which were more likely to have developed a photosynthetic capacity. The presence of light during measurement did not significantly increase the internal pO_2_ profile in buds grown in the presence (DL) or absence (D) of light (Fig. S6A), although we observed that a small increase in pO_2_ in the peripheral region of the bud (depth 0-200 µm) in the DL condition (>15 kPa *cf* D <10 kPa; Fig. S6B).

As previously observed, the spatial variance among the pO_2_ data was considerable (Fig. S5), which we interpret to be due to the considerable tissue heterogeneity within and between biological replicates, confounding the resolution of differences in oxygenation due to treatment effects. In order to establish this hypothesis, µCT were performed on individual buds after assaying the internal pO_2_ profile (168h D and DL). In the µCT the electrode path was clearly visible, enabling us to overlap the pO_2_ profile with the images (Fig. 4). As the profile supported our hypothesis, a line probe, reporting the intensity of the pixels, was used to estimate the density of tissue just above and below the electrode pathway. The pO_2_ profile correlated well with the internal structure of the bud, and particularly that the sudden decline in pO_2_ as the probe entered bract structures, while the pO_2_ of the regions of the trichome hairs was elevated (Fig. 4). This analysis also clearly illustrated that the pO_2_ of the meristematic core of the bud (between 2000-2400 µm depth) was elevated following the initiation of bud burst.

**Figure 4.**
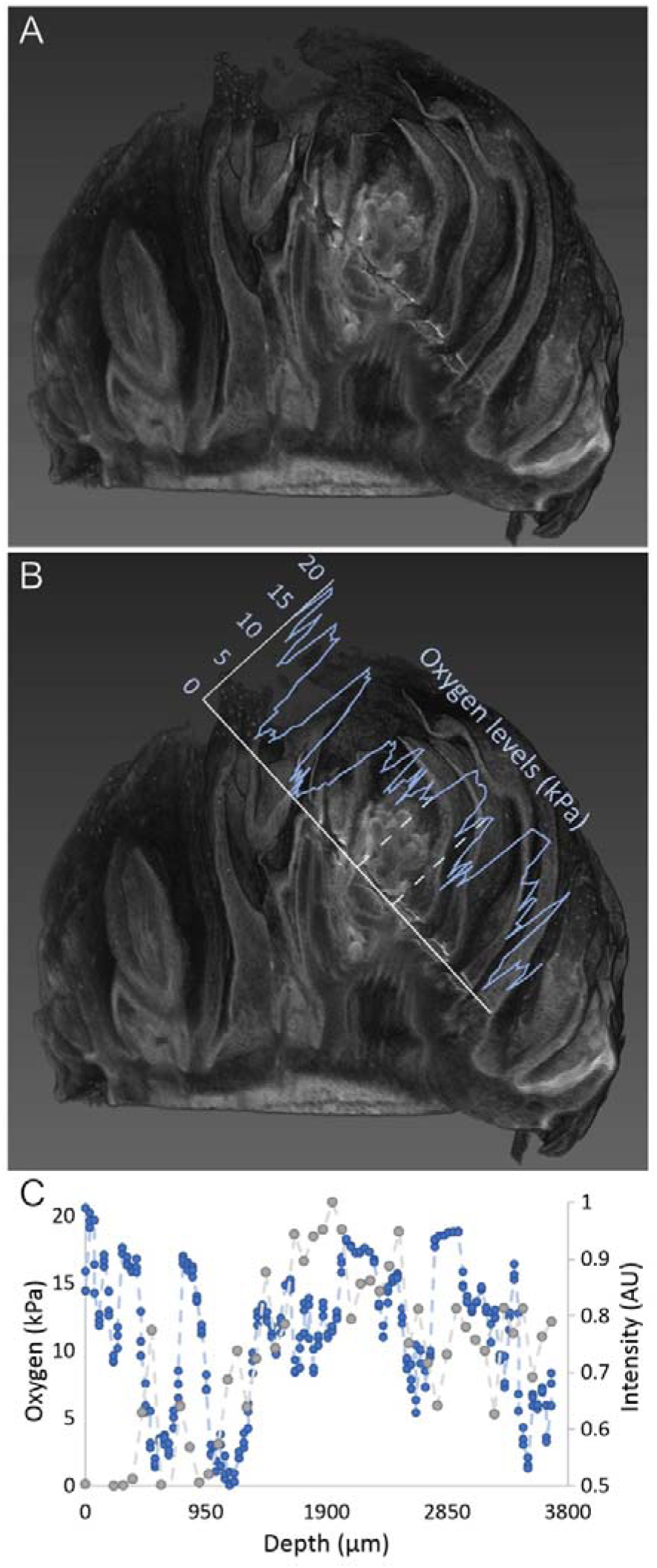
Internal oxygen profiles and micro-computed tomography of a grapevine bud. (**A**) 3D structure of the bud showing the electrode path. (**B**) O_2_ profile graph overlapped with the path of the electrode. (**c**) Internal O_2_ profile overlapped with the intensity of the signal determined by a probe line over the path in the µCT. To exemplify this figure represents the analysis of one bud (corresponding to D1 at 168h). A total of 12 buds used for internal O_2_ were scanned by µCT including 0h, 168h D and 168h DL.

Exploring the structural context further, we then determined the tissue porosity and moisture content, two variables known to affect O_2_ diffusion, of the scales, the trichomes and the green tissue of grapevine buds (Fig. 5A). These variables were first determined by weight and volume data. The greatest porosity (% gas spaces per unit tissue volume) was found in the trichomes, 77%, followed by 30% for the outer scales and 12% for the green tissue. Humidity (% water per unit tissue weight) was the lowest for the trichomes (14%), 18% for the scales and 41% for the green tissue. We further investigated the porosity analyzing the structural data obtained by µCT, being able to also evaluate the porosity of the base of the bud, the trichome, bracts and outer scales (Fig. 5B). The values were 85-88% for the trichomes, 37-38% for the outer scale, 5-6% for the bracts, and 8-10% for the base, at 0h and 168h (Fig. 5C). Overall the results from the structural images correlated well with the other results. However, the differences between the tissues were enhanced. No statistically significant differences were observed between 0h and 168h. Despite of the higher moisture and lower porosity of the inner scales, the structure of the bud and the high porosity observed (Fig. 5C) ensures that the meristem is well aerated in mature buds. On the other hand, the O_2_ profiles (Fig. 4, S5) suggests that the vascular tissue also contributes to the O_2_ concentration within the bud and not only the atmospheric O_2_ should be considered. Finally, the relatively high porosity of the outer scale determined by both methodologies suggests it is a weak barrier for oxygen.

**Figure 5.**
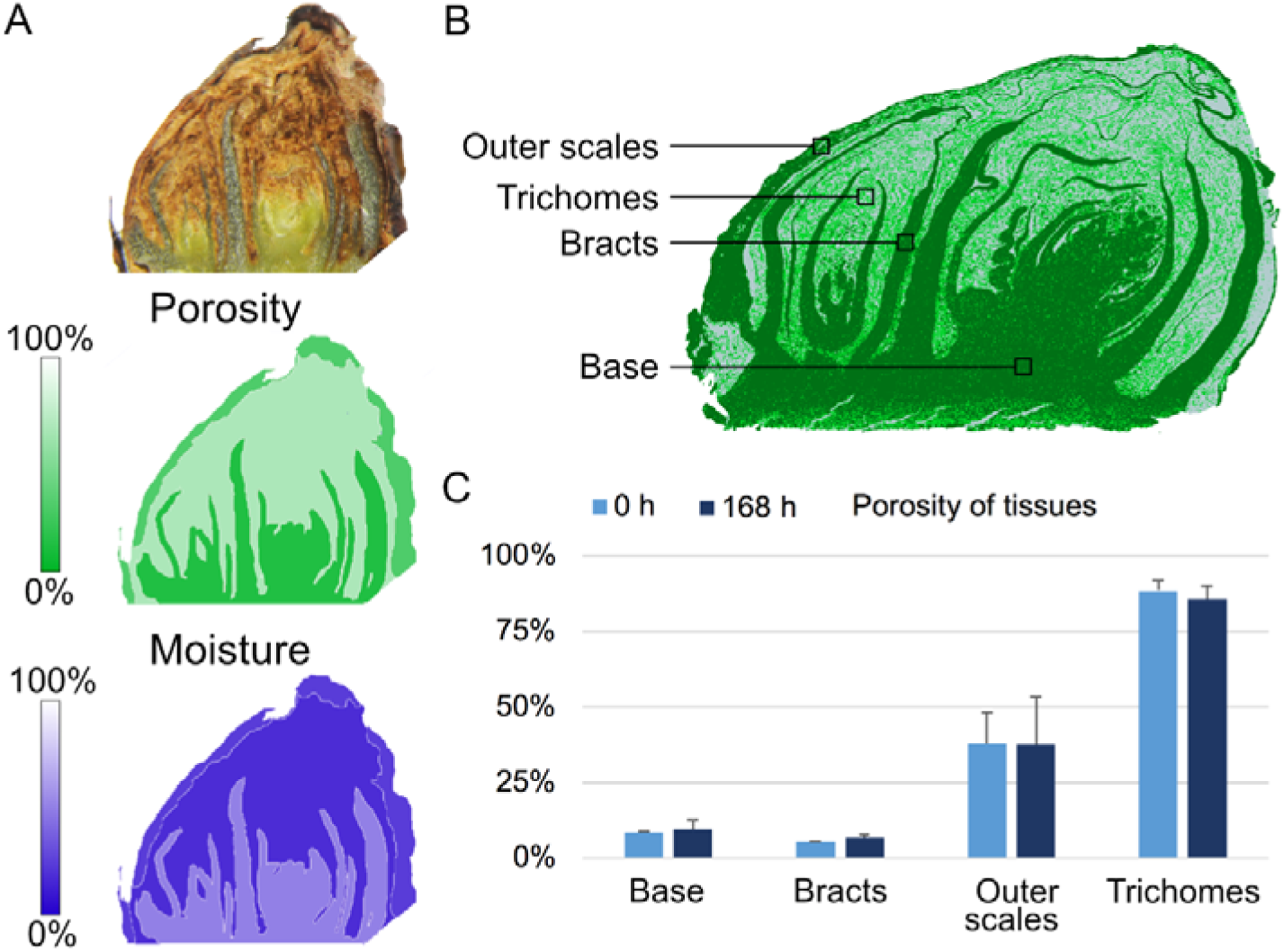
Porosity and moisture of grapevine buds. A. At the top, a sectioned bud is shown to depict the 3 evaluated parts of the bud, the green tissue, the trichomes and the outer scales. The porosity and the moisture as represented as percentage and were determined by tissue density and weight respectively. B. Section of a µCT representing the 4 type of tissues evaluated to calculate the porosity by pixel analysis. C. Comparison of porosity determined by pixel analysis at 0h and 168h.

Together these data clearly demonstrate that the pO_2_ in the bud is spatially regulated, and that despite the porosity constraints, the meristematic core of the bud (primary meristem) is preferentially oxygenated during bud burst. This oxygenation does not apparently evolve from *in situ* photosynthesis.

### Determination of the molecular exclusion size in buds

Generally speaking, the isolation of the bud from the cane during dormancy has been long considered. However, there are no data reporting quantifiable measures of this isolation, such as, the molecular exclusion size of buds. Considering that CsI was transported in 0h buds and that in grapevine dormant buds rhodamine green was shown to be transported (Jones *et al.*, 2000), the information of acid fuchsin and eosin Y can be used to precisely determine the pore size of the apoplast in grapevine prior to bud burst (Fig. S7). In order to do this, the molecules of rhodamine green, acid fuchsin and eosin Y were computationally modelled in the associated (neutral) and dissociated forms. The molecular volume, determined by the electron density of the optimised structures, was greater for eosin Y than for the other molecules (Fig. 6A). Indeed, the molecular volume (Fig. S7) positively correlated with the molecular weight of the molecules, being the lowest for rhodamine green (Fig. S7). The charges and the electrostatic potential of these molecules were also evaluated to understand if their different mobilities are due to different physical-chemical properties. In the associated form, the charges and dipole moment of rhodamine green and eosin Y are quite similar, with the partial charges homogenously distributed through the molecules (Fig. 6B). In the case of neutral acid fuchsin, more extreme partial charges were localised towards the phosphate groups, however the dipole moment was also small due to the symmetry of the molecule (Fig. 6A). Similar observations were found in the dissociated forms, with the regards to the fact that the anionic form of eosin Y had a much greater dipole moment (Fig. 6A). The fact that acid fuchsin showed more extreme partial charges would contribute to the interaction with the water molecules.

**Figure 6.**
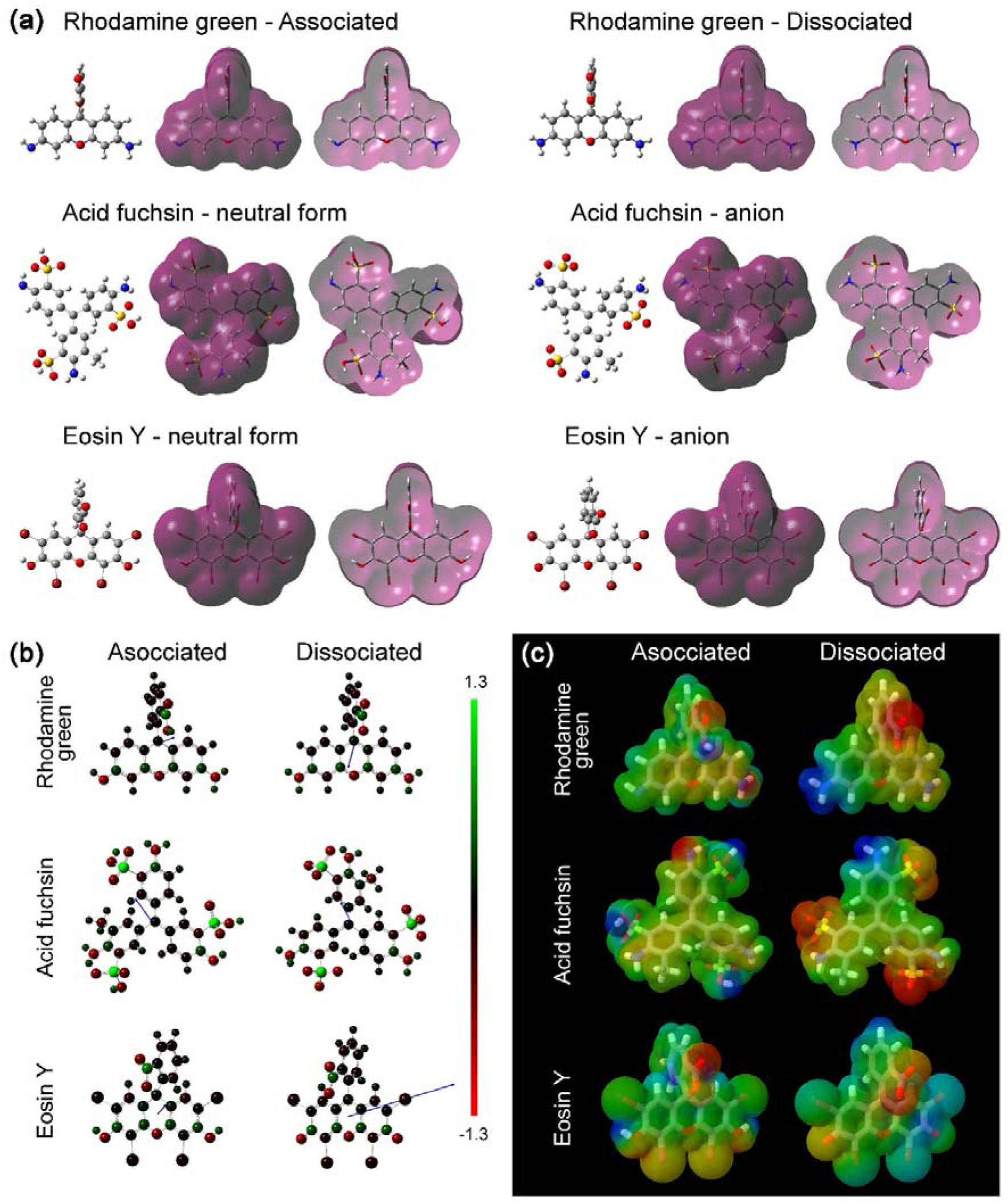
Chemical modelling of rhodamine green, acid fuchsin and eosin Y in their associated and dissociated forms. (**A**) Optimised structure and electron density. Electron density was generated using an isovalue of 0.0004. (**B**) Charges and dipole moments. The colours indicate the charge and the blue arrows indicate the dipole moment. (**C**) Electrostatic potentials. The red indicates electronegative zones whereas the blue electropositive zones.

More extreme electrostatic potentials were found in rhodamine green and acid fuchsin than in eosin Y (Fig. 6C), indicating that these two molecules are more susceptible to react with different nucleophiles and electrophiles than eosin Y. Since acid fuchsin and eosin Y are both anions (charge −2) and did not show contrasting physico-chemical properties, we can attribute the differences in mobility observed (Fig. 3B,C) to their difference in size. Hence, to better estimate the required size to transport the molecules through the plant, the solvent accessible surface for both molecules in the ionic form was modelled (Fig. 7), and the lowest area that they required to be transported was estimated. Our results demonstrated that eosin Y requires a bigger pore area to pass through (Fig. 7). Considering that these soluble molecules will require at least one layer of water hydration shell (~3.5 Å, Laage *et al*., 2017), the diameter of the pore should be increased by 0.7 nm. Hence, the 1.5 nm estimated length of these molecules would be increased to ~2.2 nm when having one water layer. As we demonstrated that these dyes cannot enter to the bud prior to bud burst, but considering that rhodamine green can, we conclude that the molecular exclusion size at the base of the bud is about 2.1 nm.

**Figure 7.**
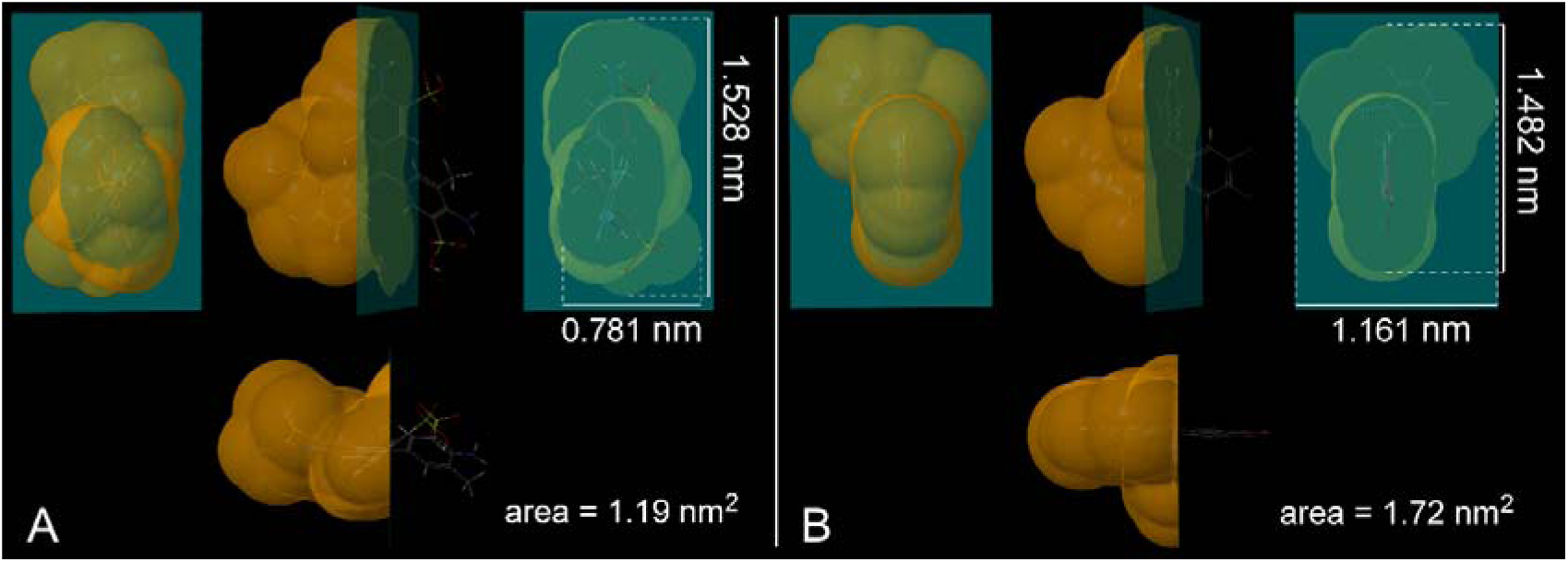
Solvent accessible surfaces for ionic forms of acid fuchsin and eosin Y. Anionic acid fuchsin. (**A**) and anionic eosin Y (**B**). A lateral, frontal and top view of each molecule is reported and the rough area of a pore that they need to pass through.

## Discussion

An increasing number of studies have added to the literature on signalling and transcriptional changes in bud quiescence, dormancy and outgrowth/ burst. Here we have returned to more classical physiological experiments to establish an improved platform of knowledge to which the gene expression and signalling studies can relate, including non-cell autonomous signals such as proteins, mRNAs and miRNAs, and the role of molecular oxygen.

### The apoplast pore size of quiescent grapevine buds restricts passive transport to water and small molecules

In the weeks prior to bud burst in woody perennials, xylem pressure builds to extreme levels and becomes enriched in phytohormones and simple sugars (Skene and Antcliff, 1972; Sperry *et al.*, 1987). It has been shown that the symplast is gated during this transition, in a manner dependent on temperature-and gibberellic acid (GA)-regulated glucanases (Rinne *et al.*, 2011). However, given the extreme pressure in the xylem, the apoplast could be a considerable pathway for long range signals from the mother vine/tree, which trigger bud burst. Experiments here show that the aperture of the apoplast would constrain transport to small molecules such as simple sugars, and may even restrict glycosylated phytohormones.

The apoplastic transport is limited by the pores formed in the cellulose/hemicellulose fiber structure. Depending on the tissue, the limiting diameter of these pores can vary. In hair cells of *Raphanus sativus* roots and fibres of *Gossypium hirsutum*, the pore size was as small as 3.5 to 3.8 nm, while in the parenchyma cells of the leaves of *Xanthium strumarium* and *Commelina communis* they were determined to be between 4.5 to 5.2 nm (Carpita *et al.*, 1979). In citrus leaves, the size exclusion limits to move through the cell wall and into the phloem was estimated to be between 4.5 to 5.4 nm (Etxeberria *et al.*, 2016). These values are much higher than the 2.1 nm diameter pore determined here in buds, suggesting that quiescent perennials buds are quite well-isolated. As a previous report demonstrated that rhodamine green, which is slightly smaller than acid fuchsin (Fig. S7), is able to pass via the apoplast into the bud (Jones *et al.*, 2000), smaller molecules should also. This would include mono- and di-saccharides, such as glucose and sucrose, and phytohormones, such as ethylene, salicylic acid, abscisic acid (ABA), jasmonic acid and GA. Xylem turgor pressure builds considerably prior to bud burst and seasonal dynamics in cytokinin (CK), sucrose and hexose content of the xylem, and transport to the bud have been correlated with bud burst in a number of woody perennials (Skene and Antcliff, 1972; Sperry *et al.*, 1987; Maurel *et al.*, 2004; Bonhomme *et al.*, 2010). For example, apical buds of the acrotonic species walnut became capable of passive and active transport of sucrose just prior to bud burst, and the transport was highly sensitive to temperature (Bonhomme *et al.*, 2010). Cell wall invertase (CWI) may play a key role in mediating the transport of sugars through the apoplast, since sucrose mobility will be more limited than for hexoses. The CWI has demonstrated functions in sink strength and regulating developmental transitions and shoot architecture (Heyer *et al.*, 2004). Indeed, CWI activity was induced shortly before bud break in buds of peach trees (Maurel *et al.*, 2004), and gene expression of a vacuolar invertase was rapidly induced at the onset of bud burst in grapevine, together with sucrose synthases (Meitha *et al*., 2018). Bonhomme *et al*. (2010) showed that by early spring the sugars were transported from the bark and xylem to the bud, at least one month prior to bud burst. Our results in excised buds collected in early spring, indicate that the *de novo* transport of sugars and phytohormones from the cane is not the trigger of bud burst. Rather, that the increase in temperature within the bud is important to enable catabolism and secondary-active transport of sugars within the bud, resulting in an increase in sink strength, which triggers a commitment to bud burst.

The fact that the pore size of the apoplastic connection is smaller in buds than in roots and leaves of other plants suggests that the isolation of the bud is key to enable desiccation and metabolic quiescence, and to avoid premature rehydration of the bud, which may result in precocious bud burst. A relevant comparison can be made to the seed coat (testa), which is known to protect the plant embryo and can restrict the transport of molecules small as 2.8 ± 1.2 nm (Kurepa *et al.*, 2010). When the seeds are imbibed, the mucilage of the epidermal cells is hydrated, rupturing the cell wall (Kurepa *et al.*, 2010). Similarly, we observed that morphological changes to bring about an increase in the pore size and allow the transport of these dyes must occur during the early stages of bud burst. Our evidence suggests that pore size increases to more than 2.1 nm over the course of bud burst to allow the diffusion of eosin Y. Importantly, here we have used computational chemistry to model the structure of the dyes at the quantum level, rather than a broadly estimated size of the molecule, as commonly used in studies reporting transport of nano-molecules. This information itself is highly valuable given that these dyes are widely employed across the life sciences to evaluate vascular transport.

### Light accelerates the reactivation of vascular tissue between the cane and the bud

Light is an absolute requirement for bud burst (outgrowth) in many species, including rosa spp. and pea (Leduc *et al.*, 2014). Here we showed that grapevine buds (post-dormant) have a facultative requirement for light, which accelerates vascular development and the rate of bud burst, but not the final proportion of buds burst. This effect was not due to a more developed state of the buds grown in DL condition, in fact, less developed DL buds showed apoplastic connectivity when more developed D buds did not (Fig. 3C, S1A,B and S2B). Our data and that of Bonhomme et al. (2010) strongly indicate that the xylem, rather than phloem being the source of metabolites for the emerging bud. In this case, re-activation of phloem as a consequence of bud burst (Esau, 1948) is secondary and not primarily related to bud burst. The local hydrolysis of polysaccharides in the cell wall should be responsible for the increase of porosity during early bud burst. In fact, upon inspecting gene expression data from a previous study (grapevine buds grown 72h and 144h in D and DL condition, data available at NCBI BioProject PRJNA327467, http://www.ncbi.nlm.nih.gov/bioproject/327467; Signorelli et al., 2018), we found that three genes coding for CELLULOSE SYNTHASE (*VIT_02s0025g01910*, *VIT_02s0025g01980* and *VIT_12s0059g00960*) and one coding for a CELLULASE (*VIT_ 19s0014g02870*) were downregulated by light, indicating a reduced metabolism of cellulose in light. Furthermore, a gene coding for EXPANSIN was upregulated by light. EXPANSINs are believe to play important roles in meristem functions, participating in the cell wall loosening (Cosgrove, 2000).

Light quality also affects bud burst in rose, in particular white, blue and red lights promote bud burst, while FR light does not, and the bud itself plays an important role in light perception (Girault *et al.*, 2008; Abidi *et al.*, 2013). In grapevine, we recently provided evidence suggesting that CRY photoreceptors play a role in promoting a photomorphogenic response, which points to the importance of blue light (Signorelli *et al.*, 2018). We are not aware of any detailed studies of the light quality responses of grapevine buds.

### The meristematic zone of buds is preferentially oxygenated during bud burst

Several studies, particularly in the presence of HC, have identified patterns that implicate the development of oxidative stress- and hypoxic response-syndromes, which precede the activation of glycolysis, the pentose phosphate pathway and fermentation, as fundamental events that enable bud burst (Ophir *et al.*, 2009; Vergara *et al.*, 2012). More recently, we observed that even in the absence of HC or other stress, tissue oxygenation and the expression of conserved hypoxia-responsive genes is acute and highly regulated during the first 24h at growth-permissive temperatures (Meitha *et al.*, 2018). Thereafter gene expression became more responsive to light and energy cues (Signorelli *et al*., 2018). Our earlier data also suggested tissue-specific oxygenation of the meristematic core (primary bud) after 24h, however the resolution of these data was not as clear as previously seen in other organs such as roots and fruit. The use of μCT data to map the path of the pO_2_ electrode here provides irrefutable evidence that the meristematic core is oxygenated during bud burst. The porosity data suggest that atmospheric pO_2_ (*ca*. 21 kPa) should be sufficient to oxygenate at least the peripheral regions and tissues of the bud. A recent study demonstrated that lenticels in the pedicel (stalk) of grapevine berries were a functional source of oxygen during ripening, linking genetic differences in this capacity to cell death and berry disorders (Xiao *et al.*, 2018). If the oxygenation of the meristem also requires oxygen from the vascular tissue, lenticels on the surface of the cane adjoining the bud may play a similar role during bud burst, however that these pathways may be occluded until bud burst has commenced and apoplastic pathways develop.

### Signals to initiate bud burst are perceived within the bud

Dormant or post-dormant perennial buds are thought to be uncoupled from apical dominance, as there is no growing shoot and the symplastic and apoplastic connections are gated. As no vascular changes were observed within the bud (Fig. 3A), and those changes observed between the bud and the cane were largely subsequent to bud burst (Fig. 3B,C, S1 and S2), we evaluated whether buds excised from the cane would be competent to establish bud burst. Our results suggested that the primary perception and signal cascade to initiate bud burst arises within the bud and are independent of *de novo* transport of phytohormones or other mobile elements from the cane. In fact, very recently, the induction of *in situ* catabolism of ABA was demonstrated to be essential for bud break in grapevine (Zheng *et al.*, 2018). Moreover, the authors showed that the transgenic expression of the ABA catabolic enzyme VvA8H-CYP707A4 resulted in enhancement of bud break in grapevine, and reduced apical dominance (Zheng *et al.*, 2018). Also recently, a transcriptomic study on *Prunus mume* suggested that low temperature results in the up-regulation of C-repeat binding factor genes, which directly promotes six dormancy associated MADS-box genes to establish dormancy. After prolonged period of cold and the subsequent rise of temperature, a decrease of the expression of these families of genes induce GA-signalling, repressing FLOWERING LOCUS T, and enabling bud burst (Zhang *et al.*, 2018).

Our data are also consistent with conclusions from other perenials plants, such as poplar, whereby the *FLOWERING LOCUS T1* (*FT1*) and *CONSTANS* (*CO*) homologues are induced within the bud by chilling, presumably synthesised in the embryonic leaves (Rinne *et al.*, 2011). Subsequent to transition to growth-permissive temperatures and long days, triggering the onset of bud burst, *FT1* expression was repressed while *CO* was elevated further. These effects were shown to be dependent on GA, and the activation of glucan hydrolases to resume symplastic communication with the cane (Rinne *et al*., 2011). Our recent transcriptome data in bursting grapevine buds also highlighted the elevated expression of GA signalling genes and those involved in core meristem functions within the first 24h transition from 4 °C to 20 °C, irrespective of light (Meitha *et al*., 2018). By contrast, at later stages, CK signalling was the prominent phytohormone signature, and demonstrably light-dependent (Signorelli *et al*., 2018). Although light and particularly photoperiod dependencies vary between poplar and grapevine, these data strong support previous conclusions that temperature, rather than light is the primary trigger for post-dormant bud burst in perennials.

Together these data suggest that bud burst in grapevine is initiated within the bud, following sufficient hydration and activation of internal ABA- and GA-dependent pathways, but independent of macromolecule transfer from the corpus. This enables an increase in metabolic activity, creating a sink and potentially signals for the resumption of vascular development.

### Outer scales of grapevine vine buds may play a role in controlling bud burst

In the present work, we showed that removing the outer scales of grapevine buds accelerates bud burst (Fig. 2A). As the effect of removing the scales was independent of light and oxygen availability (Fig. 2), we propose that the scale may facilitate biochemical repression of growth, and that HC in particular, and light more gradually, attenuate the repressor or promote its degradation. This is consistent with the biology of some seeds, where for example in legume seeds, the seed coat supplies the zygote with ABA (Smýkal *et al.*, 2014), and in Arabidopsis, a thick cutin layer surrounding the endosperm participates in GA-and ABA-dependent regulation of germination (De Giorgi *et al.*, 2015). Genetic studies showed that lines deficient in cutin biosynthesis were unable to block expansion of endosperm cells under low GA conditions. To date, studies on the regulatory role of glucans and cutins has focussed on vascular and symplastic conductance. Nevertheless, evidence from other studies in buds are consistent with the GA-dependent glucanase activity in regulating dormancy (Rinne *et al*., 2011), and this may extend to a function of a suberin layer surrounding grapevine buds, either within the outer scale or as a cicatrix-like layer between the base of the dead scale and the living bud. The primary candidate inhibitor is ABA, which was shown to inhibit dormancy release in grapevine (Zheng *et al.*, 2015, 2018). In a similar way we speculate that the bud scale could provide ABA to the bud. In some agreement, Iwasaski and Weaver (1977) showed that in grapevine bud scales, ABA rapidly increased during storage at 0 °C, reaching a maximum between the second and fourth week, and then gradually declining to control values at 12 weeks. Similarly, when HC was applied to the buds, the ABA levels rapidly increased in the scale but declined to control levels within two weeks (Iwasaki and Weaver, 1977).

## Conclusions

We conclude that the perception of environmental triggers to initiate bud burst arise within the bud, triggering metabolic and signalling activity *in situ*, which creates a sink and potentially basipetal signals for the resumption of vascular development. We also conclude that the pO_2_ patterns of buds, observed previously and here, correlated with the internal structure of the bud, in a way that the lower pO_2_ content is observed in the bract structures, whereas the pO_2_ of the regions of the trichome hairs and the meristematic core is elevated rapidly once bud burst is initiated. This explains the variance in pO_2_ within and between buds seen in this and previous studies. Given the increasingly important context of pO_2_ in plants and the use of oxygen-sensitive electrodes, these data should serve as caution to ensure the exact path is known. We also conclude that the growth repressor role of the outer scale of grapevine buds is not due to a simple physical function such as light or oxygen barrier. Finally, the smaller pore size of the apoplastic milieu of grapevine buds with respect to other plant organs allow us to conclude that the conductance of the apoplast is regulated during quiescence, although the mechanisms remain unexplored.

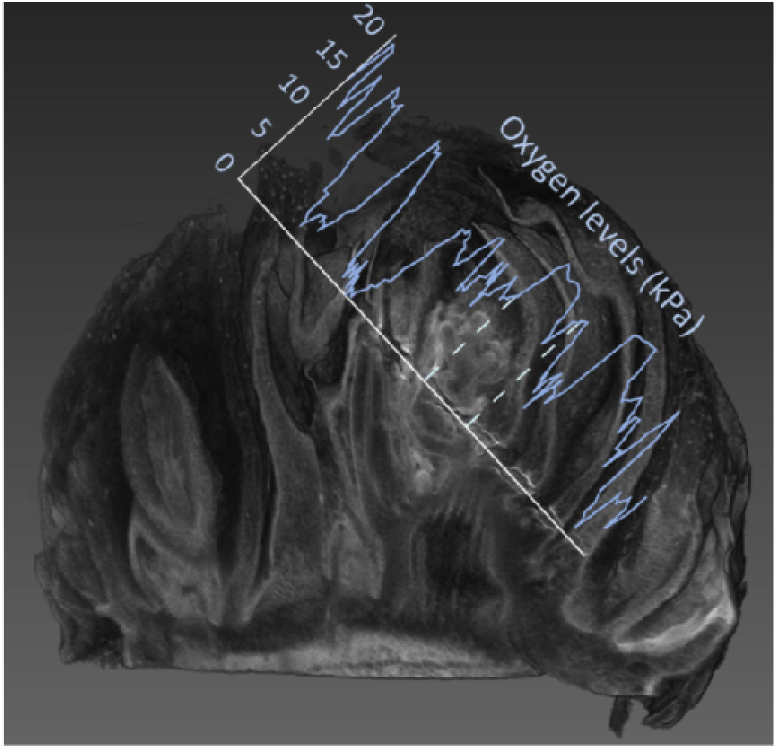

## Supplementary data

**Fig. S1:** Apoplastic connectivity in July buds at 0, 48, 96, 168, 216, 264, and 432h in D and DL.

**Fig. S2:** Apoplastic connectivity in August buds at 168, 216, and 264 in D and DL.

**Fig. S3.** Growth of isolated buds on agar.

**Fig. S4:** Apoplastic connectivity at different incubation times in 0h peeled buds and bursting buds to evaluate the effect of bud transpiration.

**Fig. S5:** Effect of light (growth condition) on internal O_2_ pressures at 0, 48, 96, and 168h in D and DL.

**Fig. S6:** Effect of light (assay condition) on measurements of internal O_2_ pressures.

**Fig. S7:** Schematic comparison of Eosin Y, Acid fuchsin and Rhodamine structures and molecular weights, to understand the logic employed to determine the molecular exclusion size.

**Movie 1:** Uptake of Iodine by grapevine bud at 168h.

## Acknowledgments

We thank Dr. Laura E. Coitiño (UdelaR, Uruguay) for her generous help in setting up the computational facilities at the School of Agronomy (UdelaR) and providing the software to perform the computational modelling. The authors acknowledge the facilities and the scientific and technical assistance of the Australian Microscopy & Microanalysis Research Facility at the Centre for Microscopy, Characterisation & Analysis, UWA, a facility funded by the University, State and Commonwealth Governments. This research was funded by Australian Research Council grants to MJC (LP0990355, DP150103211).

## Author contributions

MJC and SS conceived the study. SS carried out biological experiments, performed all physiological analysis and molecular modelling, JS, ZW, and PV performed and assisted with µCT, DH assisted with respiration measurements. SS analysed the experiments. JAC and MJC advised on experimental design. SS and MJC wrote the manuscript. All authors approved the manuscript.

## References

Abidi F, Girault T, Douillet O, Guillemain G, Sintes G, Laffaire M, Ahmed H Ben, Smiti S, Huché-Thélier L, Leduc N. 2013. Blue light effects on rose photosynthesis and photomorphogenesis. Plant Biology 15, 67–74.

Alleweldt G, Hofacker W. 1975. Influence of environmental factors on bud burst, flowering, fertility and shoot growth of vines. Vitis 14, 103–115.

Aloni R, Peterson CA. 1991. Seasonal changes in callose levels and fluorescein translocation in the phloem of Vitis vinifera L. IAWA Journal 12, 223–234.

Aloni R, Peterson C a. 1997. Auxin promotes dormancy callose removal from the phloem ofMagnolia kobus and callose accumulation and earlywood vessel differentiation inQuercus robur. Journal of Plant Research 110, 37–44.

Aloni R, Raviv A, Peterson C. 1991. The role of auxin in the removal of dormancy callose and resumption of phloem activity in Vitis vinifera. Canadian Journal of Botany 69, 1825–1832.

Beauvieux R, Wenden B, Dirlewanger E. 2018. Bud Dormancy in Perennial Fruit Tree Species: A Pivotal Role for Oxidative Cues. Frontiers in Plant Science 9, 1–13.

Bonhomme M, Peuch M, Ameglio T, Rageau R, Guilliot A, Decourteix M, Alves G, Sakr S, Lacointe A. 2010. Carbohydrate uptake from xylem vessels and its distribution among stem tissues and buds in walnut (Juglans regia L.). Tree Physiology 30, 89–102.

Carpita N, Sabularse D, Montezinos D, Delmer DP. 1979. Determination of the Pore Size of Cell Walls of Living Plant Cells. Science 205, 1144–1147.

Cartechini A, Palliotti A. 1995. Effect of shading on vine morphology and productivity and leaf gas exchange characteristics in grapevines in the field. American Journal of Enology and Viticulture 46, 227–234.

Considine MJ, Considine JA. 2016. On the language and physiology of dormancy and quiescence in plants. Journal of Experimental Botany 67, 3189–3203.

Cosgrove DJ. 2000. Loosening of plant cell walls by expansins. Nature 407, 321–326.

Esau K. 1948. Phloem structure in the grapevine and its seasonal changes. Hilgardia 18, 217–296.

Etxeberria E, Gonzalez P, Bhattacharya P, Sharma P, Ke PC. 2016. Determining the size exclusion for nanoparticles in citrus leaves. HortScience 51, 732–737.

Frisch MJ, Trucks GW, Schlegel HB, et al. 2009. Gaussian 09.

De Giorgi J, Piskurewicz U, Loubery S, Utz-Pugin A, Bailly C, Mène-Saffrané L, Lopez-Molina L. 2015. An Endosperm-Associated Cuticle Is Required for Arabidopsis Seed Viability, Dormancy and Early Control of Germination. PLoS genetics 11, e1005708.

Girault T, Bergougnoux V, Combes D, Viemont JD, Leduc N. 2008. Light controls shoot meristem organogenic activity and leaf primordia growth during bud burst in Rosa sp. Plant, Cell and Environment 31, 1534–1544.

Heyer AG, Raap M, Schroeer B, Marty B, Willmitzer L. 2004. Cell wall invertase expression at the apical meristem alters floral, architectural, and reproductive traits in Arabidopsis thaliana. Plant Journal 39, 161–169.

Iwasaki K, Weaver RJ. 1977. Effects of chilling, calcium cyanamide, and bud scale removal on bud break, rooting, and inhibitor content of buds of ‘Zinfandel’ grape (Vitis vinifera L.).

Journal of the American Society for Horticultural Science 102, 584–587.

Jones KS, McKersie BD, Paroschy J. 2000. Prevention of ice propagation by permeability barriers in bud axes of Vitis vinifera. Canadian Journal of Botany-Revue Canadienne De Botanique 78, 3–9.

Kurepa J, Paunesku T, Vogt S, Arora H, Rabatic BM, Lu J, Wanzer MB, Woloschak GE, Smalle JA. 2010. Uptake and distribution of ultra-small anatase TiO2 alizarin red s nanoconjugates in Arabidopsis thaliana. Nano Letters 10, 2296–2302.

Laage D, Elsaesser T, Hynes JT. 2017. Water Dynamics in the Hydration Shells of Biomolecules. Chemical Reviews 117, 10694–10725.

Lavee S, May P. 1997. Dormancy of grapevine buds - facts and speculation. Australian Journal of Grape and Wine Research 3, 31–46.

Leduc N, Roman H, Barbier F, Péron T, Huché-Thélier L, Lothier J, Demotes-Mainard S, Sakr S. 2014. Light Signaling in Bud Outgrowth and Branching in Plants. Plants 3, 223–250.

Maurel K, Leite GB, Bonhomme M, Guilliot A, Rageau R, Pétel G, Sakr S. 2004. Trophic control of bud break in peach (Prunus persica) trees: A possible role of hexoses. Tree Physiology 24, 579–588.

May P, Clingeleffer P, Brien C. 1976. Sultana (Vitis vinifera L.) canes and their exposure to light. Vitis 14, 278–288.

Meitha K, Agudelo-Romero P, Signorelli S, Gibbs DJ, Considine JA, Foyer CH, Considine MJ. 2018. Developmental control of hypoxia during bud burst in grapevine. Plant Cell and Environment 41, 1154–1170.

Meitha K, Konnerup D, Colmer TD, Considine JA, Foyer CH, Considine MJ. 2015. Spatio-temporal relief from hypoxia and production of reactive oxygen species during bud burst in grapevine (Vitis vinifera). Annals of Botany 116, 703–711.

Michailidis M, Karagiannis E, Tanou G, Sarrou E, Adamakis ID, Karamanoli K, Martens S, Molassiotis A. 2018. Metabolic mechanisms underpinning vegetative bud dormancy release and shoot development in sweet cherry. Environmental and Experimental Botany 155, 1–11.

Nicolas WJ, Grison MS, Trépout S, Gaston A, Fouché M, Cordelières FP, Oparka K, Tilsner J, Brocard L, Bayer EM. 2017. Architecture and permeability of post-cytokinesis plasmodesmata lacking cytoplasmic sleeves. Nature Plants 3.

Ophir R, Pang X, Halaly T, Venkateswari J, Lavee S, Galbraith D, Or E. 2009. Gene-expression profiling of grape bud response to two alternative dormancy-release stimuli expose possible links between impaired mitochondrial activity, hypoxia, ethylene-ABA interplay and cell enlargement. Plant Molecular Biology 71, 403–423.

Or E, Vilozny I, Eyal Y, Ogrodovitch A. 2000. The transduction of the signal for grape bud dormancy breaking induced by hydrogen cyanamide may involve the SNF-like protein kinase GDBRPK. Plant Molecular Biology 43, 483–494.

Or E, Vilozny I, Fennell A, Eyal Y, Ogrodovitch A. 2002. Dormancy in grape buds: Isolation and characterization of catalase cDNA and analysis of its expression following chemical induction of bud dormancy release. Plant Science 162, 121–130.

Parada F, Noriega X, Dantas D, Bressan-Smith R, Pérez FJ. 2016. Differences in respiration between dormant and non-dormant buds suggest the involvement of ABA in the development of endodormancy in grapevines. Journal of Plant Physiology 201, 71–78.

Paul LK, Rinne PLH, Van der Schoot C. 2014. Shoot meristems of deciduous woody perennials: Self-organization and morphogenetic transitions. Current Opinion in Plant Biology 17, 85–95.

Petrie PR, Clingeleffer PR. 2005. Effects of temperature and light (before and after budburst) on inflorescence morphology and flower number of Chardonnay grapevines (Vitis vinifera L.). Australian Journal of Grape and Wine Research 11, 59–65.

Possingham J V. 2004. On the growing of grapevines in the tropics. Acta Horticulturae.39–44.

Rinne PLH, Welling A, Vahala J, Ripel L, Ruonala R, Kangasjärvi J, van Der Schoot C. 2011. Chilling of Dormant Buds Hyperinduces FLOWERING LOCUS T and Recruits GA-Inducible 1,3-b-Glucanases to Reopen Signal Conduits and Release Dormancy in Populus. The Plant Cell 23, 130–146.

Sánchez LA, Dokoozlian NK. 2005. Bud microclimate and fruitfulness in Vitis vinifera L. American Journal of Enology and Viticulture 56, 319–329.

Signorelli S, Agudelo-Romero P, Considine MJ, Foyer CH. 2018. Roles for light, energy and oxygen in the fate of quiescent axillary buds. Plant Physiology 176, 1171–1181.

Skene KGM. 1967. Gibberellin-like substances in root exudate of Vitis vinifera. Planta 74, 250–262.

Skene KGM, Antcliff AJ. 1972. A comparative study of cytokinin levels in bleeding sap of Vitis vinifera (L.) and the two grapevine rootstocks, salt creek and 1613. Journal of Experimental Botany 23, 282–293.

Smýkal P, Vernoud V, Blair MW, Soukup A, Thompson RD. 2014. The role of the testa during development and in establishment of dormancy of the legume seed. Frontiers in Plant Science 5, 351.

Sperry JS, Holbrook NM, Zimmermann MH, Tyree MT. 1987. Spring filling of xylem vessels in wild grapevine. Plant physiology 83, 414–417.

The Jmol Team. 2007. Jmol: an open-source Java viewer for chemical structures in 3D. Jmolsourceforgenet.

Tilsner J, Nicolas W, Rosado A, Bayer EM. 2016. Staying Tight: Plasmodesmal Membrane Contact Sites and the Control of Cell-to-Cell Connectivity in Plants. Annual Review of Plant Biology 67, 337–364.

Vergara R, Rubio S, Pérez FJ. 2012. Hypoxia and hydrogen cyanamide induce bud-break and up-regulate hypoxic responsive genes (HRG) and VvFT in grapevine-buds. Plant Molecular Biology 79, 171–178.

Wang Z, Verboven P, Nicolai B. 2017. Contrast-enhanced 3D micro-CT of plant tissues using different impregnation techniques. Plant Methods 13, 1–16.

Xiao Z, Rogiers SY, Sadras VO, Tyerman SD. 2018. Hypoxia in grape berries: The role of seed respiration and lenticels on the berry pedicel and the possible link to cell death. Journal of Experimental Botany 69, 2071–2083.

Zhang Z, Zhuo X, Zhao K, Zheng T, Han Y, Yuan C, Zhang Q. 2018. Transcriptome Profiles Reveal the Crucial Roles of Hormone and Sugar in the Bud Dormancy of Prunus mume. Scientific Reports 8, 1–15.

Zheng C, Acheampong AK, Shi Z, et al. 2018. Abscisic acid catabolism enhances dormancy release of grapevine buds. Plant Cell and Environment 41, 2490–2503.

Zheng C, Halaly T, Acheampong AK, Takebayashi Y, Jikumaru Y, Kamiya Y, Or E. 2015. Abscisic acid (ABA) regulates grape bud dormancy, and dormancy release stimuli may act through modification of ABA metabolism. Journal of Experimental Botany 66, 1527–1542.

